# Linking medicinal cannabis to autotaxin-lysophosphatic acid signaling

**DOI:** 10.1101/2022.07.01.498446

**Authors:** Mathias C. Eymery, Andrew A. McCarthy, Jens Hausmann

## Abstract

Autotaxin is primarily known for the formation of lysophosphatidic acid (LPA) from lysophosphatidylcholine (LPC). LPA is an important signaling phospholipid that can bind to six G-protein coupled receptors (LPA_1-6_). The ATX-LPA signaling axis is a critical component in many physiological and pathophysiological conditions. Here, we describe a potent inhibition of Δ^9^-*trans*-tetrahydrocannabinol (THC), the main psychoactive compound of medicinal cannabis and related cannabinoids, on the catalysis of two isoforms of ATX with nanomolar EC_50_ values. Furthermore, we decipher the binding interface of ATX to THC, and its derivative 9(R)-Δ6a,10a-THC (6a10aTHC), by X-ray crystallography. Cellular experiments confirm this inhibitory effect, revealing a significant reduction of internalized LPA_1_ in the presence of THC with simultaneous ATX and LPC stimulation. Our results establish a functional interaction of THC with autotaxin-lysophosphatidic acid signaling and highlight novel aspects of medicinal cannabis therapy.

## Introduction

Autotaxin (ATX; ENPP2) is an extracellular glycoprotein, which hydrolyzes lysophosphatidylcholine (LPC) into lysophosphatidic acid (LPA) by cleaving off the choline head group^1, 2, 3^. LPA is a multifunctional bioactive lipid mediator with six designated G-protein coupled receptors (LPA_1-6_)^4^, forming together with ATX the ATX-LPA signaling axis. ATX is the main producer of LPA in serum, which has been demonstrated by a heterozygous *Enpp2* (ATX) knock-out mouse model. These mice show only 50% of the normal LPA levels in serum^5^. It is widely accepted that the ATX-LPA signaling axis is of crucial importance for lipid homeostasis in humans^6^. ATX is present in almost every body fluid and essential for murine embryonic vessel formation, which highlights its importance for life^3, 7, 8^. Thus, the ATX-LPA axis is linked to numerous physiological and pathological processes, such as vascular and neuronal development, neuropathic pain, fibrosis, immune-mediated diseases including rheumatoid arthritis, multiple sclerosis and atherosclerosis and cancer^3^. In fact *Enpp2* (ATX) is among the top 40 most up-regulated genes in metastatic breast cancer^9^, while ATX-LPA signaling is positively correlated with the invasive and metastatic potential of several cancers including melanoma, breast, ovarian, thyroid, renal cell, lung, neuroblastoma, hepatocellular carcinoma and glioblastoma multiforme^10^.

ATX consists of four domains, two repetitive N-terminal somatomedin B-like domains (SMB1 and SMB2), followed by the catalytic phosphodiesterase domain (PDE) and an inactive nuclease domain (Nuc)^11, 12^. The active site of ATX constitutes a bimetallo zinc coordination center and the active site nucleophile, Thr209 in rodents^11^. A nearby hydrophobic pocket, which extends into the PDE domain, accommodates the lipid substrates aliphatic chain; additionally there is an allosteric tunnel that is formed between the SMB2 and PDE domains, where an oxysterol and bile acids bind^11,13^.

The gene product of ATX can exist in at least three different isoforms, which are ATX-α, ATX-β and ATX-γ, as a result of an alternative splicing event^14^. ATX-α is characterized by a polybasic insertion of 52 amino acids in the PDE domain, when compared to the canonical plasma isoform ATX-β. ATX-α can bind to heparin and cell surface heparan sulfate proteoglycans^15^, while ATX-β has been shown to bind to β_1_ and β_3_ subunits of integrins^11, 16^. ATX-γ is the so-called “brain-specific” isoform^17^ and has been implicated with neuronal disorders, like multiple sclerosis, depression, Alzheimer’s Disease and neuropathic pain^3^.

Another important signaling system is the well-established endocannabinoid system^18^, with its two cannabinoid receptors, the cannabinoid receptor type 1 (CB_1_) and type 2 (CB_2_)^19, 20^. The human CB_1_ is primarily expressed in the central nervous system and also present in the peripheral nervous system and testis^19^, while the CB_2_ receptor is mainly expressed in the immune system^20^. The identification of their endogenous ligands, anandamide (AEA)^21^ and 2-arachidonoylglycerol (2-AG)^22^, which were detected in samples of the brain and intestine, and shown to activate CB_1_ and CB_2_ with high affinity and efficacy, were subsequently described as endocannabinoids^23,18^. The endocannabinoid system can be further expanded to the endocannabinoidome, a much wider complex network of promiscuous mediators overlapping with other signaling pathways, including LPA and its receptors^18^. Interestingly, it has been shown that dephosphorylation of a 2-arachidonoyl species of LPA in the brain of rats leads to the formation of the endocannabinoid 2-AG^24^, a process that was later revealed to depend on lipid phosphate phosphatases^25^. Additionally, the two endocannabinoid receptors, CB_1_ and CB_2_, show an amino acid sequence identity to LPA_1-3_ of around 18-20%^26^. Moreover, a functional crosstalk between CB_1_ and LPA_1_ has been revealed, where the absence of the main cerebral receptors for LPA or endocannabinoids is able to induce a modulation on the other at the levels of both signaling and synthesis of endogenous neurotransmitters^27^.

Pharmacological manipulation of the endocannabinoid system can be achieved by medicinal cannabis. The major psychoactive cannabinoid component of medicinal cannabis from the plant *Cannabis sativa* is Δ^9^-*trans*-tetrahydrocannabinol (THC), which can bind to CB_1_ and CB_2_ in a low nanomolar regime^28^. Here, we show the potential of THC, and other cannabinoid compounds, to modulate the catalytic activity of ATX, and present results that THC can reduce ATX-mediated LPA signaling in a cellular context.

## Results and Discussion

### Inhibition of ATX by various cannabinoids

We first set out to validate the hypothesis that cannabinoids might bind ATX to modulate its catalytic function. For this we utilized various cannabinoids (Fig. 1 A) at a fixed concentration of 10 μM for each compound in our biochemical validation with LPC 18:1 (200 μM) as substrates in an end-point assay for ATX-β (Uniport: Q13822-1) and ATX-γ (Uniport: Q13822-3) (Fig. 1 B and C). The quality of our enzyme assay is confirmed with HA155, a well-documented ATX inhibitor (Fig. EV1 A)^29, 11^. The obtained IC_50_ (6 ± 0.8 nM) for HA155 is similar in our assay conditions compared to previous results^29^. Additionally, we can exclude an interference of the cannabinoid compounds in our enzymatically coupled assay, as no difference is detectable in the absence of ATX and LPC (Fig. EV1 B).

**Fig. 1:**
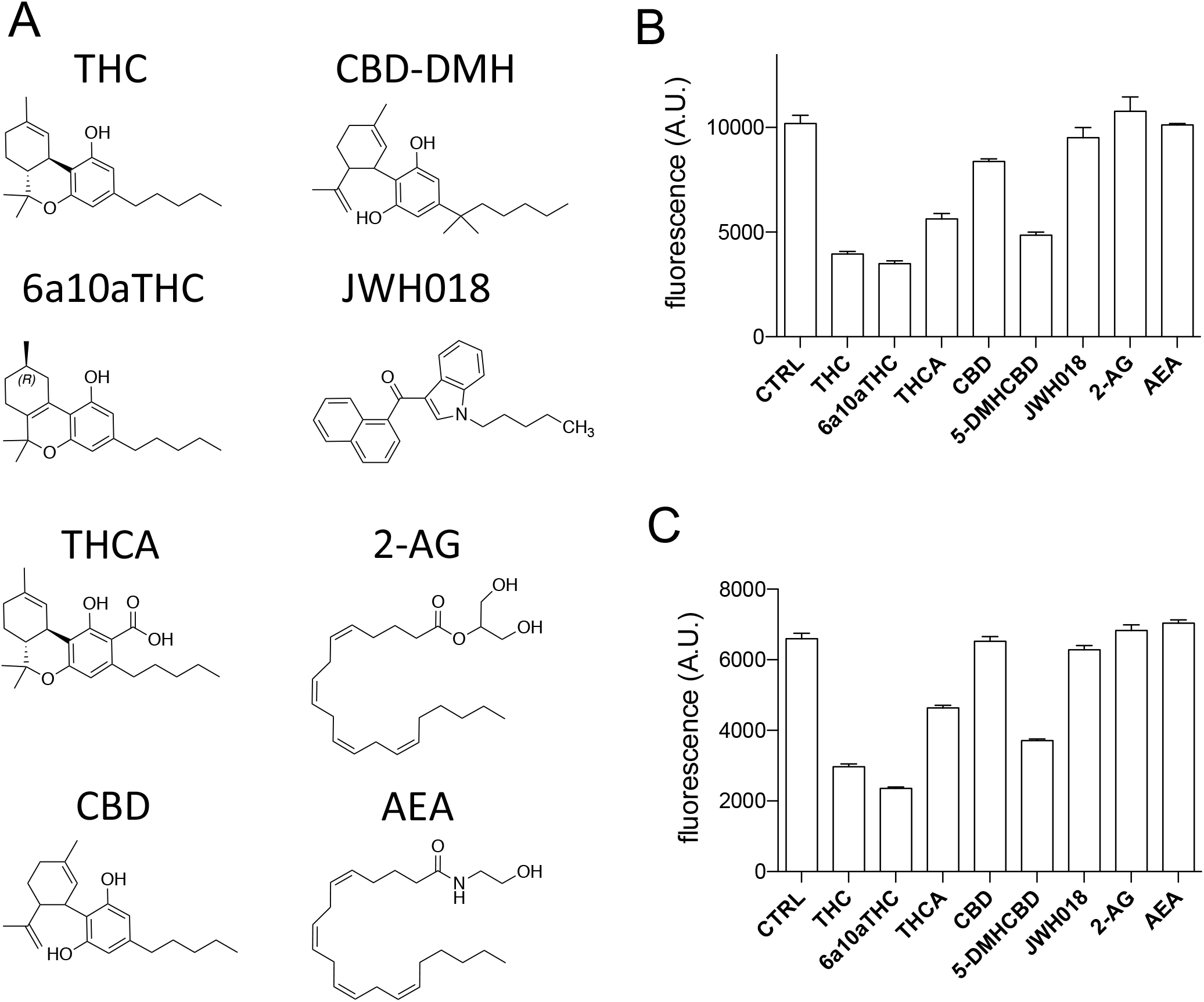
End point assays of compounds tested (A) Chemical representation of Δ^9^-*trans*-tetrahydrocannabinol (THC), Cannabidiol-dimethylheptyl (CBD-DMH), 9(R)-Δ6a,10a-THC (6a10aTHC), JWH018, Tetrahydrocannabinolic acid (THCA), 2-arachidonoylglycerol (2-AG), Cannabidiol (CBD) and Anandamide (AEA). End-point assay for (B) ATX-β and (C) ATX-γ inhibition with various cannabinoids and endocannabinoids. All error bars represent the s.e.m (n=3).

We observe a potent inhibition of THC on the catalysis of both ATX isoforms with more than 50% inhibition (Fig. 1 B and C). Furthermore, 9(R)-Δ6a,10a-THC (for simplicity referred to from here as: 6a10aTHC), a derivative of THC that differs only in the position of the double bond in the C-ring compared with THC (Fig. 1 A) is included in our cannabinoid inhibition screen. Interestingly, this minimal difference causes a further increase in the magnitude of inhibition for both ATX isoforms tested (Fig. 1 B and C). Tetrahydrocannabinolic acid (THCA) is a precursor of THC and an active component of medicinal cannabis. It is distinguishable from THC by the presence of a carboxylic group at the A-ring (Fig. 1 A). THCA also showed an inhibition of the enzymatic activity of both ATX isoforms tested. However, this inhibition is less pronounced, when compared to THC and 6a10aTHC, and did not reach a 50% inhibition magnitude in our assay conditions (Fig. 1 B and C).

The next compound we tested was cannabidiol (CBD), a non-psychoactive ingredient of medicinal cannabis. CBD is structurally different from THC by an opening of the B-ring. Interestingly, CBD showed only a weak inhibition towards ATX-β, and no observable inhibition for ATX-γ (Fig. 1 B and C). CDB-DMH is a synthetic CBD derivative, which is characterized by the addition of two methyl groups at the beginning of the aliphatic chain and an elongation with a single methyl group at the end of the carbon chain (Fig. 1 A). These structural modifications remarkably increase the magnitude of inhibition for ATX-β and ATX-γ (Fig. 1 B and C).

We also analyzed JWH018 (Fig. 1 A), which is a synthesized compound and full agonist for CB_1_ and CB_2_ with K_i_ values of 9.0 ± 8.0 nM and 2.9 ± 2.7 nM, respectively^30^. However, this artificial cannabinoid did not influence the catalytic activity of either ATX isoform (Fig. 1 B and C). To complete our picture of the modulation cannabinoids on the enzymatic activity of ATX, we also utilized the endocannabinoids 2-AG and AEA. However, both endocannabinoids did not affect the catalysis of ATX in the applied conditions (Fig. 1 B and C).

### Biochemical characterization of THC and 6a10aTHC with ATX-β and ATX-γ

We choose THC and 6a10aTHC for our detailed biochemical characterization, as these inhibitors have a maximum magnitude of inhibition of more than 50%, a cut-off criterion selected under the assay conditions used. THC works as a partial inhibitor on the catalysis of both isoforms (Fig. 2 A and B). The EC_50_ values of THC with ATX-β and LPC 18:1 as substrate is 1025 ± 138 nM, as shown in Fig. 2 A. The magnitude of inhibition is around 60%. A similar magnitude of inhibition is observed with ATX-γ, with an EC_50_ of 407 ± 67 nM for THC towards this isoform (Fig. 2 B).

**Fig. 2:**
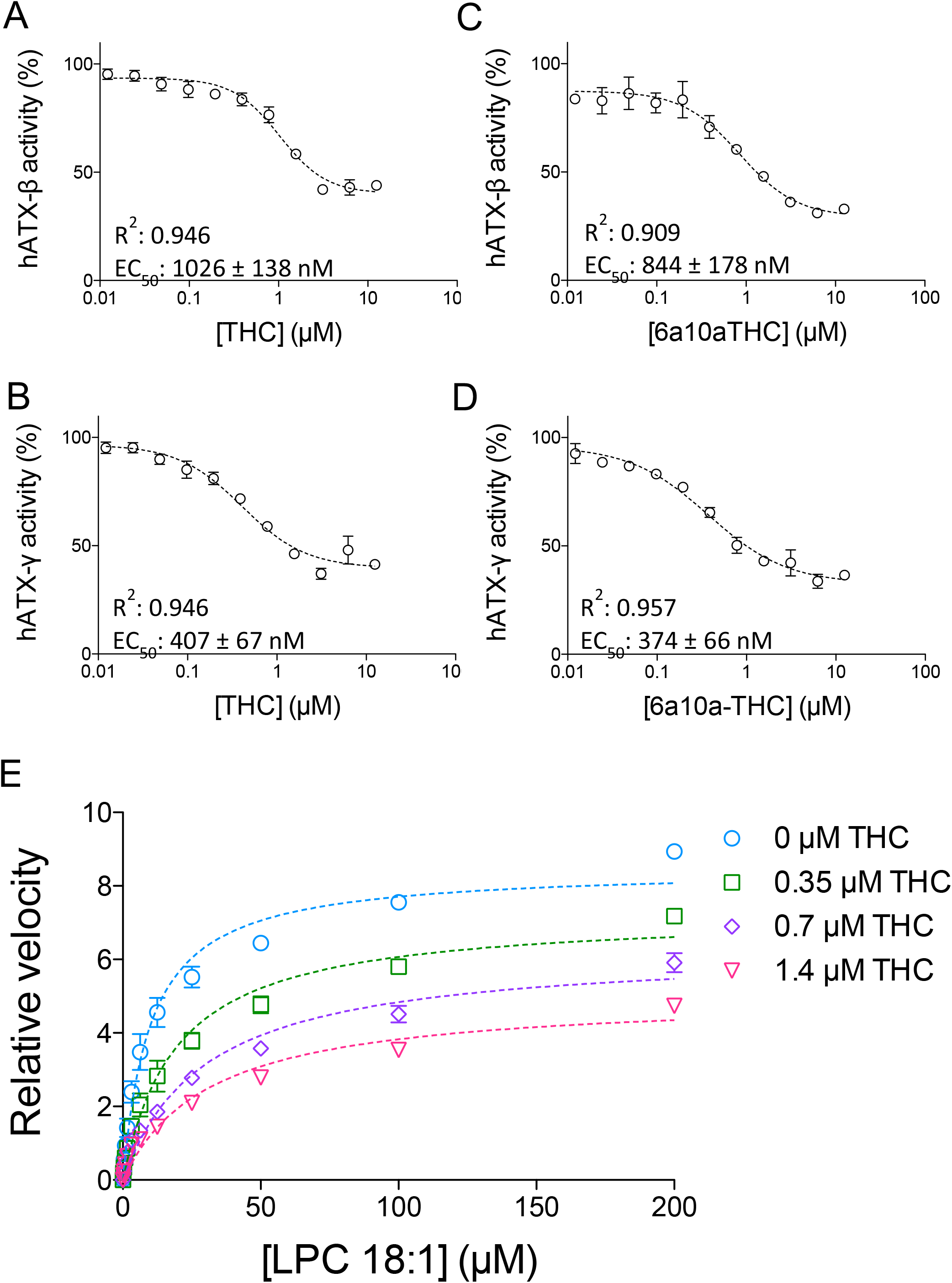
Inhibition of ATX by plant-derived THC and synthetic 6a10aTHC. Dose-response analysis of (A) ATX-β and (B) ATX-γ with THC and LPC 18:1, (C) ATX-β and (D) ATX-γ with 6a10aTHC and LPC 18:1. (E) Mode of inhibition of THC with ATX-γ indicates a mixed type inhibition. All error bars represent the s.e.m (n=3).

Next, we validate the artificial THC derivative 6a10aTHC. The EC_50_ value of 6a10aTHC for ATX-β is 844 ± 178 nM (Fig. 2 C) and thus, comparable to THC. The maximum inhibition is marginally increased and appears to be around 75%. 6a10aTHC has the highest potency towards ATX-γ with a determined EC_50_ of 374 ± 66 nM (Fig. 2 D). The magnitude of inhibition is around 70%, which is consistent with ATX-β. Overall, 6a10aTHC is the best utilized cannabinoid inhibitor for both isoforms tested with the classical substrate LPC 18:1, and also for LPC 16:0 (Fig. EV2).

Both THC and 6a10aTHC are very closely related in structure to each other, and show a similar behavior in biochemical assays. Thus, we performed a mode of inhibition analysis with THC only, in order to understand the inhibition mode of these compounds (Fig. 2 E). This analysis is carried out with 0, 0.35, 0.7 and 1.4 μM of THC with geometrically increasing concentrations of LPC 18:1. It revealed that THC functions as a mixed-type inhibitor, which is demonstrated by the decrease of V_max_ from 8.5 to 7.3, 6.3 and 5.0, respectively, and an increase in K_m_ from 10.1 to 19.6, 29.9 and 31.5 μM, respectively.

### Co-crystal structure of ATX-THC

To understand the binding interface between THC and ATX in detail, we expressed and purified the second isoform of ATX from *Rattus norvegicus* (UniProt ID: Q64610-2) and co-crystallized this formerly used ATX construct^11,13^ with THC. We determined this ATX-THC structure (PDB ID: 7P4J) to 1.8 Å resolution with an R_free_ of 23.5% (Table 1). We obtained clear residual electron density close the active site of ATX. Modelling of THC here resulted in a very good fit to this remaining electron density (Fig. 3 A). ATX binds THC at the entrance of the hydrophobic pocket with the aliphatic chain pointing into this pocket. The binding of the THC molecule is driven by hydrophobic interactions of the residues I167, F210, L213, L216, W254, F274, Y306 and V365 (Fig. 3 B), as analyzed by the PLIP server^31^. A superposition of our ATX-THC structure with the ATX-LPA 18:1 structure (PDB ID: 5DLW)^13^ shows that the THC molecule blocks binding of the LPA 18:1 aliphatic chain, while binding to the glycerol backbone and the phosphate group can still occur (Fig. 3 C).

**Table 1.** Crystallographic data collection and refinement statistics

| Crystal | ATX-THC | ATX-9(R)-Δ6a,10a-THC |
| --- | --- | --- |
| PDB identifier | 7P4J | 7P4O |
| *Data collection* |  |  |
| Wavelength (Å) | 0.976 | 1.000 |
| Space group | P1 | P1 |
| *Cell dimensions* |  |  |
| a, b, c (Å) | 53.7 61.0 63.6 | 53.8 62.4 64.4 |
| $\alpha$, $\beta$, $\gamma$ (°) | 103.2 97.4 94.2 | 103.7 98.4 93.4 |
| Resolution (Å)* | 61.2-1.8 (1.9-1.8) | 53.0-1.7 (1.75-1.7) |
| No. of reflections | 47909 (2395) | 85204 (8434) |
| R<sub>pim</sub> (%) | 5.8 (64) | 7.25 (63.8) |
| Completeness (%) |  |  |
| Spherical | 65.9 (13.1) | 94.8 (94.2) |
| Ellipsoidal | 91.6 (60.4) | - |
| Redundancy | 9.2 (7.0) | 3.5 (3.6) |
| *Refinement* |  |  |
| R<sub>work</sub> (%) | 18.69(29.1) | 17.10 (21.5) |
| R<sub>free</sub> (%) | 23.5(22.8) | 20.60 (25.3) |
| No. of atoms<sup>†</sup> | 6800 | 6818 |
| Protein+carbohydrates | 6230 | 6244 |
| Ligand+metal ions | 163 | 113 |
| Waters and other ions | 407 | 461 |
| B-factors (Å<sup>2</sup>) |  |  |
| All | 27.5 | 31.9 |
| Protein+carbohydrates | 27.0 | 31.3 |
| Ligand+metal ions | 34.3 | 42.1 |
| Water and other ions | 32.9 | 36.6 |
| R.m.s. deviations |  |  |
| Bond lengths (RMS) | 0.005 | 0.007 |
| Bond angles (RMS) | 0.86 | 0.90 |

\*Values given in parenthesis refer to reflections in the highest resolution bin. For calculation of R<sub>free</sub> 5% of all reflections were omitted from refinement. <sup>†</sup>Alternate conformations are counted as multiple atoms.

**Fig. 3:**
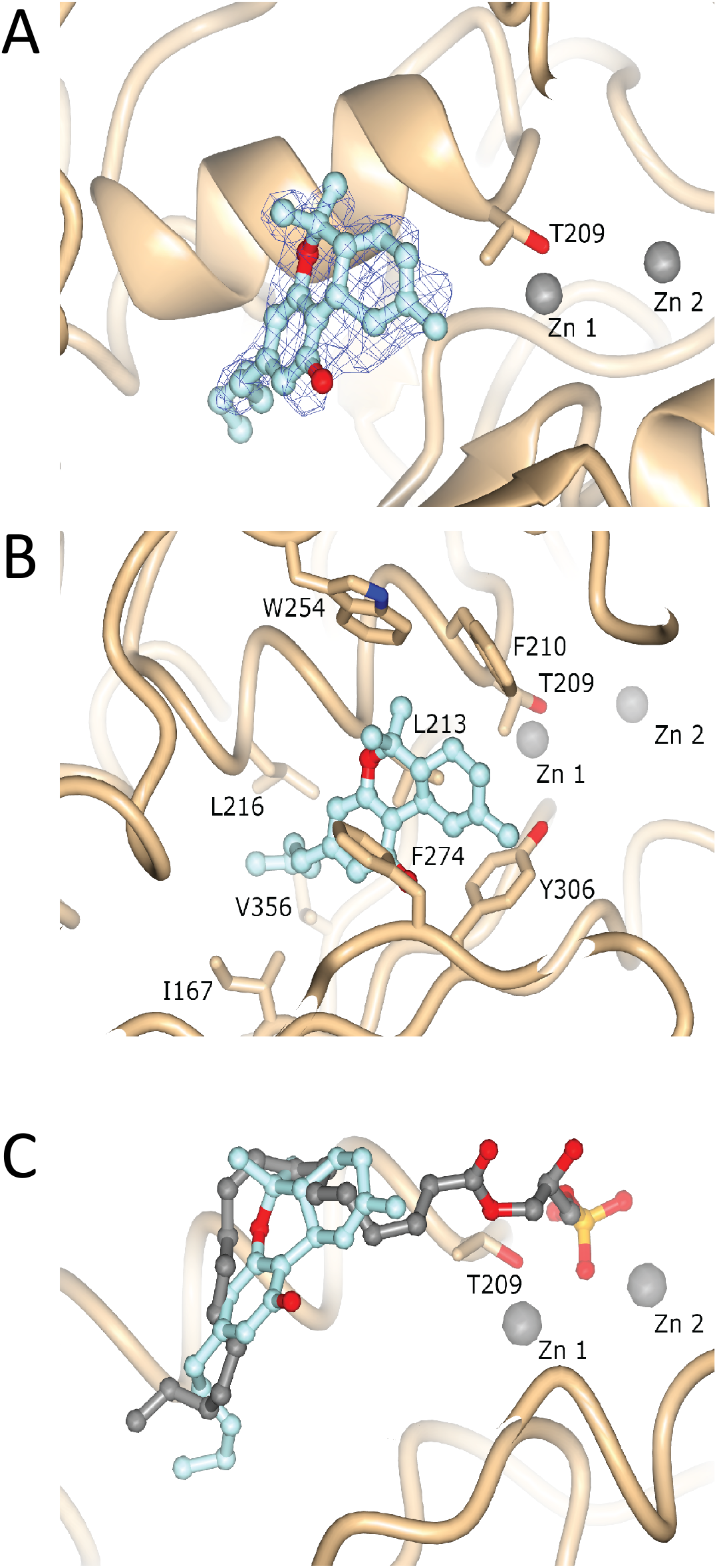
Structure and electron density of ATX-THC (A) Feature Enhanced electron density map before THC placement, contoured at 1 RMSD and represented as a blue wireframe model. (B) Molecular interactions of THC with ATX. (C) Superposition of ATX binding to THC (PDB ID: 7P4J) and LPA 18:1 (PDB ID: 5DLW).

### Co-crystal structure of ATX-6a10aTHC

We also obtained an ATX-6a10aTHC structure (PDB ID: 7P4O) to 1.7 Å resolution with an R_free_ of 20.6% (Table 1). In this ATX-6a10aTHC structure, we observed clear residual electron density, which resembles almost perfectly the 6a10aTHC ligand (Fig. 4 A). The binding of the 6a10aTHC molecule is again mainly accomplished by hydrophobic interactions of the residues I167, F210, L213, W254, F273, F274, Y306 (Fig. 4 B), as analyzed by the PLIP server^31^. Nevertheless, an additional water bridge between the carbonyl of F273 and the THC derivative can be observed (Fig. 4 C), which suggests that the binding stability of this ligand is higher compared to THC, and potentially explains the lower EC_50_ for 6a10aTHC. However, the authors are aware that a comparable water molecule also exists in the ATX-THC structure, where the distance of the THC oxygen and carbonyl oxygen of F273 is 4.5 Å, thus above the PLIP server threshold for such an interaction (Fig. EV3).

**Fig. 4:**
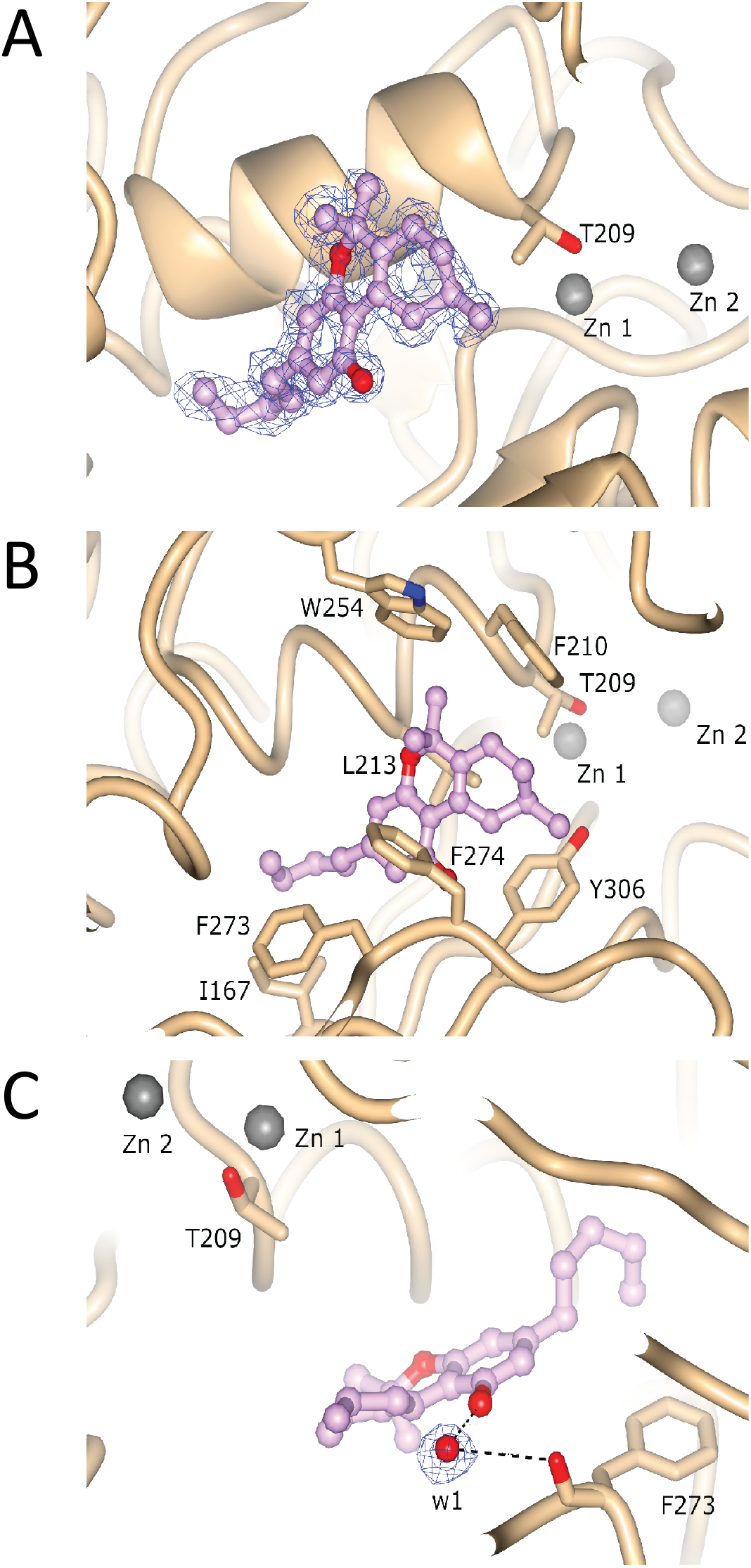
Structure and electron density of ATX-6a10aTHC. (A) Featured Enhanced electron density map before 6a10aTHC placement, contoured at 1 RMSD and represented as a blue wireframe model. (B) Molecular interactions of 6a10aTHC with ATX. (C) Bridging water molecule interaction between 6a10aTHC and carbonyl oxygen of F273.

The 6a10aTHC ligand in ATX adopts an overall similar binding position to the cannabinoid in our ATX-THC structure. However, the aliphatic chain of the ligand points in a slightly different direction when compared to THC. Also noteworthy is that the cyclohexene (C-ring) appears to adopt a different stereoisomeric configuration (Fig. EV4) due to the alternate localization of the double bond (Fig. 1 A).

### Inhibition of LPA_1_ internalization in Hela cells by THC

To validate THC can act as an inhibitor in the production of LPA and thus, ATX-LPA signaling in an cellular context, we used an agonist-induced LPA_1_ receptor internalization as a readout in cultured cell assays^32, 33, 34^. As shown in Fig. 5, stimulation of HA/LPA_1_ transfected HeLa cells with 30 nM ATX, 150 μM LPC 18:1, and 1 μM THC significantly reduced LPA_1_ internalization. This observation was only detectable in the presence of ATX and not in control conditions (Fig. EV5). This is an indirect response to blocking LPA production, which inhibits receptor activation and endocytosis, confirming a more physiological role of THC as a potent inhibitor.

**Fig. 5.**
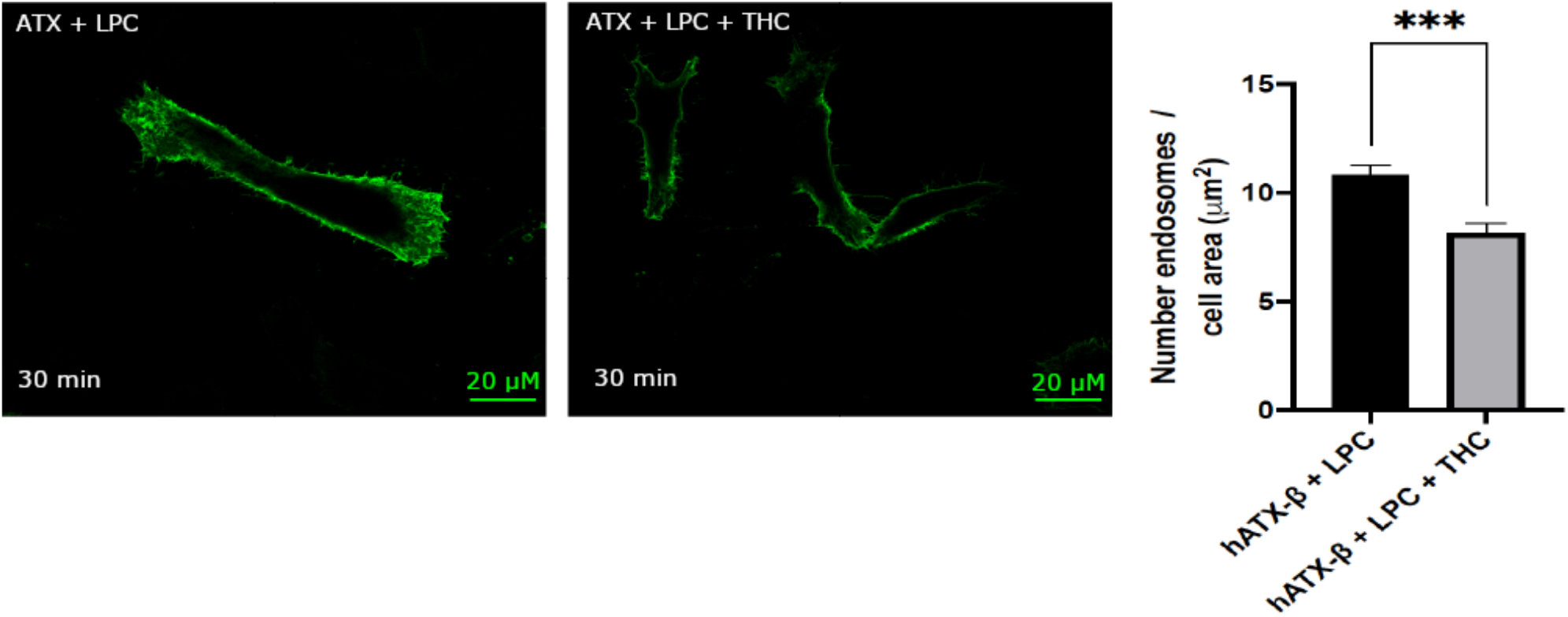
Physiological effect of THC on LPA_1_ receptor internalization. Quantification of LPA_1_ receptor internalization revealed that THC reduced the number of endosomes internalized when compared to untreated condition, paired t-test p-value = 0.0008. All error bars represent the SEM, calculated from 11 images per condition in biological triplicate experiments.

## Discussion

Medicinal cannabis has been approved as a therapeutic agent by local authorities in an increasing number of states all over the world. Even though great progress in the molecular basis of medicinal cannabis therapy has been achieved over the last decades, the pleiotropic effects have been insufficiently characterized to date. We establish here a potent *in vitro* inhibition of various cannabinoids, such as THC, on the catalysis of ATX with different substrates (LPC 16:0 and LPC 18:1) and isoforms. Based on our investigations, we provide evidence that THC can potently modulate LPA signaling.

In most studies that try to address the pharmacological aspects of medicinal cannabis, the administration has been performed via smoking. In this context, the first body fluid that encounters THC is the saliva. The mean concentration of THC in this oral fluid has been detected with up to 4167 ng/ml (13 μM) in a radioimmunoassay^35^. Interestingly, LPA is present in saliva^36^ and ATX expression can be detected in salivary gland tissue^37^, suggesting that ATX-LPA signaling may be reduced by THC *in vivo*. Furthermore, the deposition of THC in oral fluid reflects a similar time course in plasma after smoked cannabis administration^38^. Serum concentrations of THC show a wide inter-individual difference, between 59-421 ng/ml after a 49.1 mg THC dose, which equals 190 nM to 1.3 μM of THC^39^. The observed mean THC peak in this study for a 69.4 mg THC dose was 190.4 ng/ml (SD=106.8), which is in the range of the EC_50_ determined during our studies.

95-97% of THC is bound to plasma proteins and our data suggest that ATX might function as a carrier for THC in plasma. The authors extensively tried to determine the binding affinity of ATX for THC with various techniques, such as isothermal titration calorimetry and nuclear magnetic resonance, however these approaches were unsuccessful due to the hydrophobic nature of the cannabinoid ligand.

Recently, the potential use of medicinal cannabis in tissue fibrosis has been proposed^40^. In this regard, it is noteworthy to mention the role of ATX in idiopathic pulmonary fibrosis (IPF). Several inhibitors targeting ATX are under clinical investigations for their therapeutic use against IPF^41^. However, the ISABELA study (clinical phase 3 investigation) of the most advanced molecule targeting ATX, Ziritaxestat (Glpg1690) from Galapagos N.V, was discontinued due to risk-benefit concerns. It is tempting to speculate, that a full ATX inhibitor, which reduces LPA levels to almost zero, causes many systemic unwanted side effects, as the ATX-LPA signaling axis is pivotal under physiological conditions. In this context, our observation that THC is a partial inhibitor of ATX is of great interest, because this molecule is an FDA approved drug, which reduces LPA levels incompletely. Moreover, the fact that THC can cross the blood-brain-barrier makes it an attractive candidate to manipulate neuronal diseases, where the brain-specific isoform of ATX is involved.

Additionally, glaucoma is the leading cause of irreversible blindness worldwide^42^. Glaucoma is characterized by elevated intraocular pressure (IOP) levels and medicinal cannabis is being used to treat this pathology, however the therapeutic mechanism is not completely known. Interestingly, in recent years it has been discovered that aqueous humor samples of patients suffering from primary open angle glaucoma have elevated levels of ATX, LPC and LPA^43^. Moreover, pharmacological inhibition of ATX lowered IOP in rabbits^44^. Our data may explain the molecular basis for the therapeutic effect of medical cannabis in glaucoma patients, as THC could feasibly reduce the formation of LPA by inhibiting the enzymatic activity of ATX. In conclusion, our study warrants further research into the pleiotropic effects of medicinal cannabis in the context of ATX-LPA signaling, while also providing a promising starting point for such research lines. Furthermore, this work also provides a scaffold for the design of new inhibitors for further studies of the ATX-LPA signaling axis, and suggests a new way to intervene in ATX-LPA signaling-mediated pathologies with THC.

## Material and Methods

### Materials

We obtained T300 tissue culture flasks (#90301) from TPP and roller bottles (#681070) from Greiner Bio-One; DMEM medium (Gibco #12491023), Opti-MEM (Gibco #31985062); FBS (Gibco #10270106), fatty acid free FBS (Gibco #A3382101), L-Glutamine (Gibco #25030-123), POROS-20 MC column (Thermo Scientific™ #1542906), lipofectamine 3000 (Thermo Scientific™ #L3000001), Alexa Fluor™ 594 Conjugate (Invitrogen # W11262) and ; SDS precast gel (Invitrogen #XP04205BOX) from Thermo Fisher Scientific; CELLSTAR 12 well culture plates (Greiner #665180) and Fluoroshield (#F6182-20mL) from Sigma-Aldrich; Amicon ultra 15 ml 10 kDa (#UFC901008) and ultra 0.5 ml 10 kDa (#UFC501008) concentrators from Merck; Superose 6 (10/300) column (#17-5172-01) and Superdex 200 increase 3.2/300 column (#28-9909-46) from GE Healthcare; Trans-blot turbo transfer pack (#1704158) from BioRad; LPC 18:1 (#845875P), LPC 16:0 (#855675C), and LPA 18:1 (#857130C) from Avanti Polar Lipids. Choline quantification kits (#40007) from AAT Bioquest; THC (#LGCAMP1088.00-05) from LGC Standards, France; 9(R)-Δ6a,10a-THC (#33013) from Cayman. CBN (#C-046-1ML) from Cerilliant; 5-DMH-CBD (#1481) from Tocris; CBD (#HB2785) from Hellobio; NH_4_I (#AB202711) from Abcr; NaSCN (#HR2-693) from Hampton Research; InstantBlue Coomassie Protein stain (#ab119211), anti-HA tag primary antibody (#ab18181), anti-mouse antibody (#ab150113), anti-ATX antibody (#ab77104), anti-mouse HRP secondary antibody (#ab6728), and ECL substrate kit (#ab133406) from Abcam.

### Methods

#### ATX expression and purification

Recombinant ATX proteins were essentially produced as previously described^45^. HEK293-Flp-In cells were cultivated in complete DMEM medium supplemented with 10% FBS with minor differences. The cells were grown to 80-90% confluence, washed twice with preheated PBS and trypsinized for 5 min with 5 ml of trypsin. Inactivation was accomplished by adding 45 ml of complete medium. Cells were resuspended in complete medium and inoculated into roller bottles. 10 T300 flasks were used to inoculate 8 roller bottles and the cells were cultured for 4 days after transfer into 125 ml DMEM containing 10% FBS and 2 mM glutamine. The medium was then replaced by 125 ml DMEM containing 2% FBS and 2 mM glutamine. The cells were then left to express protein for 4 to 5 days before collection. Fresh expression medium was added for a further round of recombinant expression.

HEK293 medium from eight roller bottles was pooled together and the recombinant ATX proteins were purified using a POROS-20 MC column pre-loaded with Cu^2+^. Equillibration was achieved by washing with ten column volumes of buffer A (20 mM HEPES, 150 mM NaCl, pH 7.4). ATX was eluted with a linear gradient of buffer B consisting of buffer A supplemented with 500 mM imidazole. Reasonably pure fractions were pooled after SDS-PAGE analysis. The fraction volume was reduced with an Amicon ultra 15 ml 10 kDa concentrator to a volume of 500 μl. 5 mg/ml concentrated protein was injected on Superose 6 (10/300) gel filtration column using buffer A. The purity of the peak fractions was analyzed by SDS-PAGE. The recombinant protein concentration was determined by the ratio of the optical density at 260/280 nm using a Nanodrop 2000 (Thermo Fischer Scientific). The ATX construct from *Rattus Norvegicus* (UniProt ID: Q64610-2) was concentrated to 3-3.5 mg/ml using an Amicon Ultra 0.5 ml 10 kDa concentrator. *Human* ATX (UniProt ID: Q13822-1 and Q13822-3) was concentrated to 1.3 mg/ml. Purity was assessed using SDS-PAGE, Western blot and SEC analysis (Fig. EV6 A-C). For the SDS-page, 25 μg of purified protein in reducing conditions were loaded on a precast gel, run for one hour at 225 volt and imaged after InstantBlue Coomassie Protein stain after following the manufacturer’s instructions. For Western blot, proteins were transferred using Trans-Blot turbo transfer system (BioRad). Membrane staining with the primary antibody was performed overnight using an anti-ATX antibody. Anti-mouse HRP secondary antibody was incubated for one hour and detection was then performed using an ECL substrate kit. Analytical SEC was performed by injecting 25 μL of rATX on a Superdex 200 increase 3.2/300 equilibrated with a 50mM Tris-HCL (pH 8) and 150 mM NaCl buffer. hATX-β and hATX-γ activity was controlled using the choline release assay described below (Fig. EV6 D). hATX-γ activity was slightly lower than hATX-β activity, which is in accordance with published comparison of the different ATX isoforms^14^.

#### End-point assays

ATX phospholipase D activity was measured using choline release from LPC18:1 and LPC 16:0 with a choline quantification kit^46^ (Fig. EV6 D). 30 nM ATX-β or γ was incubated with 200 μM LPC 18:1 or LPC 16:0 in a final volume of 100 μl buffer, which contained 50 mM Tris-HCl (pH 8.5) and 150 mM NaCl. The experiments for determining relative inhibition for various cannabinoids were performed at 37 °C by adding 10 μM of the cannabinoid or endocannabinoid mentioned. Released choline was detected and the enzyme activity was determined by measuring fluorescence at λ_ex_/λ_em_=540/590 nm in 96-well plates, every 60 seconds for 50 min minimum using a Clariostar plate reader (BMG Labtech). Absolute values were taken at 25 min after visual inspection and the 0 min baselines were subtracted to account for compound differences. The relative inhibition values were determined using the normalize method in Graphpad Prism (GraphPad Software, Inc). Measurements have been performed in triplicate with three different protein preparations. All the compounds were controlled for interference of fluorescence and inhibition in the same assay conditions but in the absence of ATX and replacing LPC with choline.

#### Dose response assay for cannabinoids

ATX phospholipase D activity was measured using choline release from LPC18:1 and LPC 16:0 with a choline quantification kit^46^. 30 nM ATX-β or γ were incubated with 200 μM LPC 18:1 or LPC 16:0 in a final volume of 100 μl buffer, which contained 50 mM Tris-HCl (pH 8.5) and 150 mM NaCl. The experiment for determining EC_50_ for various cannabinoids were performed at 37 °C by adding the cannabinoid in a serial 2-fold dilution for each concentration. For THC and 6a10aTHC the retained 2-fold dilution started at 12.5 μM. For CBN, 5-DMH-CBD and CBD the starting concentration was 50 μM, 150 μM and 2 mM, respectively. Released choline was detected and the enzyme activity was determined by measuring fluorescence at λ_ex_/λ_em_=540/590 nm in 96-well plates, every 60 seconds for 50 min minimum using a Clariostar plate reader (BMG Labtech). Initial velocities were taken between 19-31 min after visual inspection. The EC_50_ values were determined using the non-linear regression analysis method (fit: [inhibitor] vs. response (three parameter)) in Graphpad Prism (GraphPad Software, Inc). Measurements have been performed in triplicate with three different protein preparations.

#### Crystallization, structure determination and model building

Crystallization experiments were performed at 303 K using the hanging-drop vapor diffusion method as previously published^47^. The best crystals were obtained with the *r*ATX construct (3-3.5 mg/ml) after 30 min RT preincubation with 5 mM THC or 5 mM 6a10aTHC dissolved in ethanol. 1 μl of the protein solution was then mixed with 1μl of the reservoir solution containing 18-22% (m/v) PEG3350, 0.1-0.3 M NH_4_I and 0.3 M NaSCN. All the crystals were cryoprotected with the addition of 20% (v/v) of glycerol.

X-ray data for THC and 6a10aTHC autotaxin complexes were collected at 100K on EMBL PETRA III beamline P14 and P13^48^, respectively. Crystallographic ATX-THC complex data were acquired using the Global Phasing WFs data collection workflow to maximize the completeness of the P1 dataset. Authorization to collect sample containing THC was granted by the BfArM in Germany. All data were processed with Autoproc^49^ staraniso which includes XDS^50^. Structures were determined by molecular replacement using MRage^51^ with the structure of ATX (PDB: 2XR9) as model^11^. Model building was performed using Coot^52^, phenix.refine^53^, REFMAC5^54^ and PDB-REDO^55^. Ligands were drawn with ELBOW^56^. Validation of the model was performed with phenix PDB deposition tools, using MolProbility^57^. Maps were generated using phenix refine and feature-enhanced map^58^. The crystallographic parameters and model quality indicators can be found in Table 1. Structural figures were generated using CCP4mg^59^. Structural biology applications used in this project were compiled and configured by SBGrid^60^.

#### hLPA_1_ receptor internalization assay

The hLPA_1_ receptor internalization assay was essentially performed as previously described^34^. A pRP[Exp]-Puro-CMV>HA/hLPA_1_ vector coding for full length human LPA_1_ receptor (UniProt ID: Q92633) with a human influenza hemagglutinin (HA) sequence epitope-tag at the 5'-end of the extracellular domain was designed and maxiprep plasmid DNA was produced commercially (Vector Builder). Vector quality control was done by restriction enzyme analysis and Sanger sequencing.

HeLa cells were grown on coverslips in a 12 well plate format and transfected with HA/hLPA_1_ vector in DMEM complete medium with lipofectamine 3000 using 1 μg of plasmid DNA, and 3 μL of Lipofectamine 3000 per well after complexation in 50 μL Opti-MEM, as per the manufacturer’s instructions, 48 hours prior to fixation. 8 hours prior to treatment and fixation, the cells were starved in fatty acid free DMEM to avoid hLPA_1_ activation by serum lipids. Several assays were performed in different conditions before fixation: 30 nM ATX + 150 μM LPC 18:1; 30 nM ATX + 150 μM LPC 18:1 + 1 μM THC; 1 μM THC; 1 μM LPA 18:1; Untreated (vehicle only); and untransfected: to control specificity of the antibody towards HA tagged hLPA_1_ receptor. LPC 18:1 and LPA 18:1 were dissolved in fatty acid free FBS with a final concentration in the media of 0.1%. THC was dissolved in DMSO to a final concentration in the media of 0.025% (v/v) DMSO.

Fixation was carried out by adding paraformaldehyde directly into the media to a final concentration of 3%, and incubating at 37° C for 10 min. Cells were washed 3 times in PBS and membranes were labelled using Wheat Germ Agglutinin, and Alexa Fluor™ 594 Conjugate for 10 min at 5 μg/mL in PBS, as per the manufacturer’s instructions. Cells were washed three times in PBS, permeabilized using 0.2% tween for 10 min, washed in PBS, and finally blocked with 10% goat serum for 30 min. HA tag was labelled using an anti-HA tag primary antibody at 1/200 dilution in 10% FBS for 1 hour at room temperature followed by PBS wash and secondary staining was done with an anti-mouse antibody, 30 min incubation at 1/500 dilution in 10% FBS. Cells were washed three times and mounted using Fluoroshield mounting. Imaging was performed using a Leica SP5 (X 63 objective). Endosome quantification was done using Fiji analyze particle tools after image thresholding. The number of counted endosomes were normalized over the measuring area to calculate the density per μm^2^. Statistical analysis was performed using paired t-test over 11 images for each condition of ATX-THC-LPC and ATX-LPC in biological triplicate.

## Acknowledgements

The authors are grateful for the initial *in silico* work of Dr. Ulrike Uhrig of the Chemical Biology Core Facility, EMBL, who demonstrated a potential inhibition of THC towards ATX, which was fundamental to apply for an initial THC license. The authors are also grateful to Dr. Corinna Gorny for her support to obtain a license for Dronabinol experiments at EMBL Germany under BfArM authorization. We are thankful for the kind gift of ATX expressing cell lines from the Perrakis lab in Amsterdam. We thank beamline staff from the EMBL-Hamburg beamlines at the PETRA III storage ring (DESY, Hamburg, Germany) for beamtime (proposal number MX-661). In particular we thank Drs. G. Bourenkov, as well as G. Bricogne and R. Fogh at Global Phasing Limited who provided valuable assistance with multi-orientation data collection strategies on P14. The authors also wish to express their gratitude to the Eukaryotic Expression Facility in Grenoble for infrastructure access, especially Alice Aubert and Martin Pelosse for excellent technical support and fruitful discussions. Lastly, we thank L. Gutierrez for her technical support with biochemical assays. MCE has been funded by the EMBL International PhD program. AAM has been funded by EMBL. JH was supported by a fellowship from the EMBL Interdisciplinary Postdoc (EI3POD) programme under Marie Skłodowska-Curie Actions COFUND (grant number: 664726).

## Expanded Figures

**Fig. EV1.**
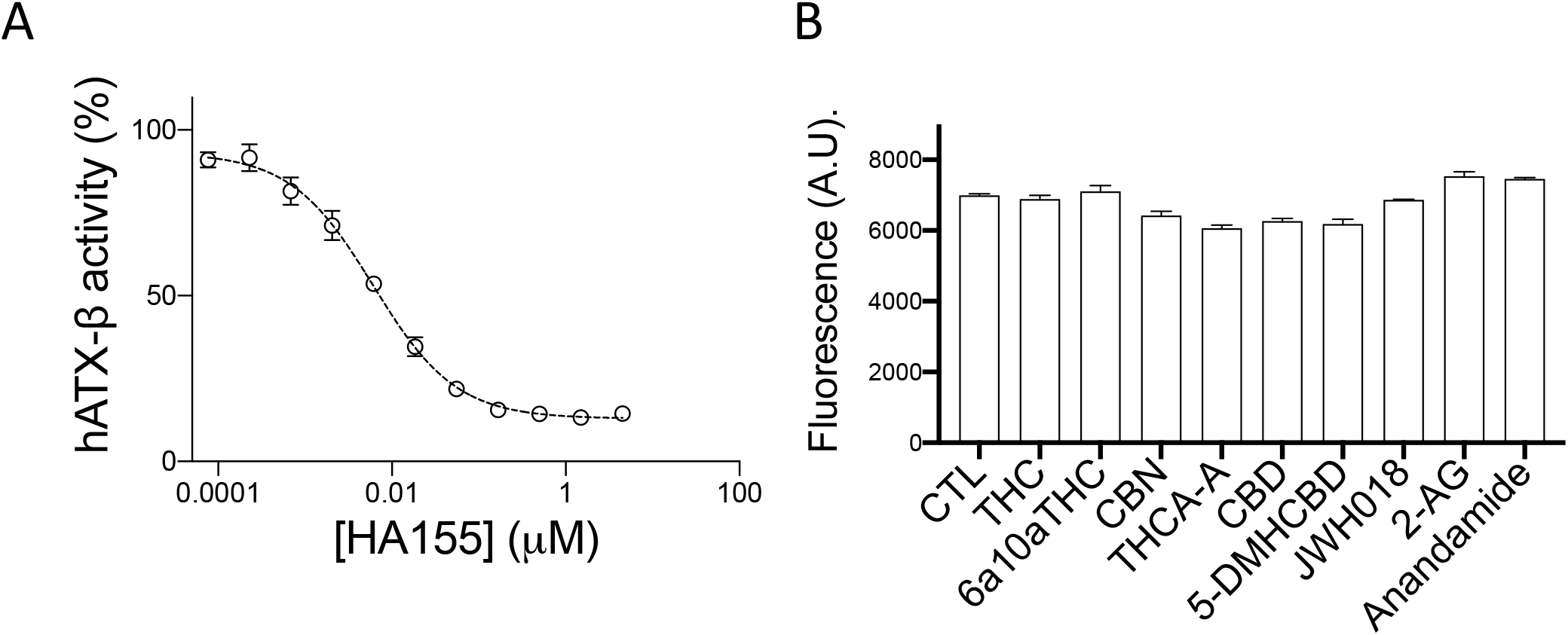
Enzymatic assay controls (A) Dose-response analysis of HA155 with LPC 18:1 and ATX-β showing an IC50 of 6 ± 0.8 nM. (B) Control assay in absence of ATX and replacement of LPC with choline. No interference with the assay was found with the various cannabinoids and endocannabinoids tested. All error bars represent the s.e.m (n=3).

**Fig. EV2.**
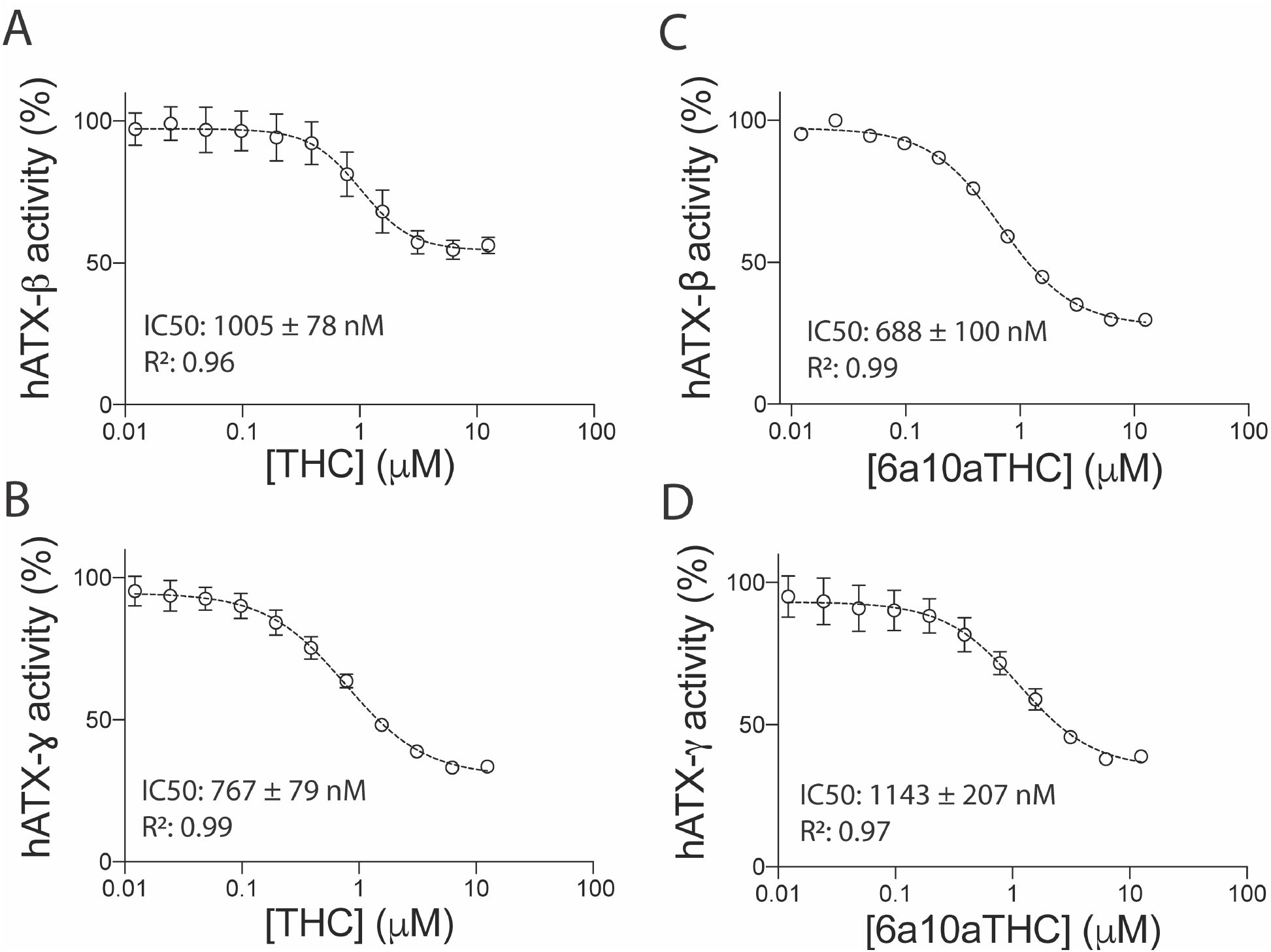
Inhibition of ATX by plant-derived THC and synthetic 6a10aTHC. Dose-response analysis of (A) ATX-β and (B) ATX-γ with THC and LPC 16:0. Dose-response analysis of (C) ATX-β and (D) ATX-γ with 6a10aTHC and LPC 16:0. All error bars represent the s.e.m (n=3).

**Fig. EV3.**
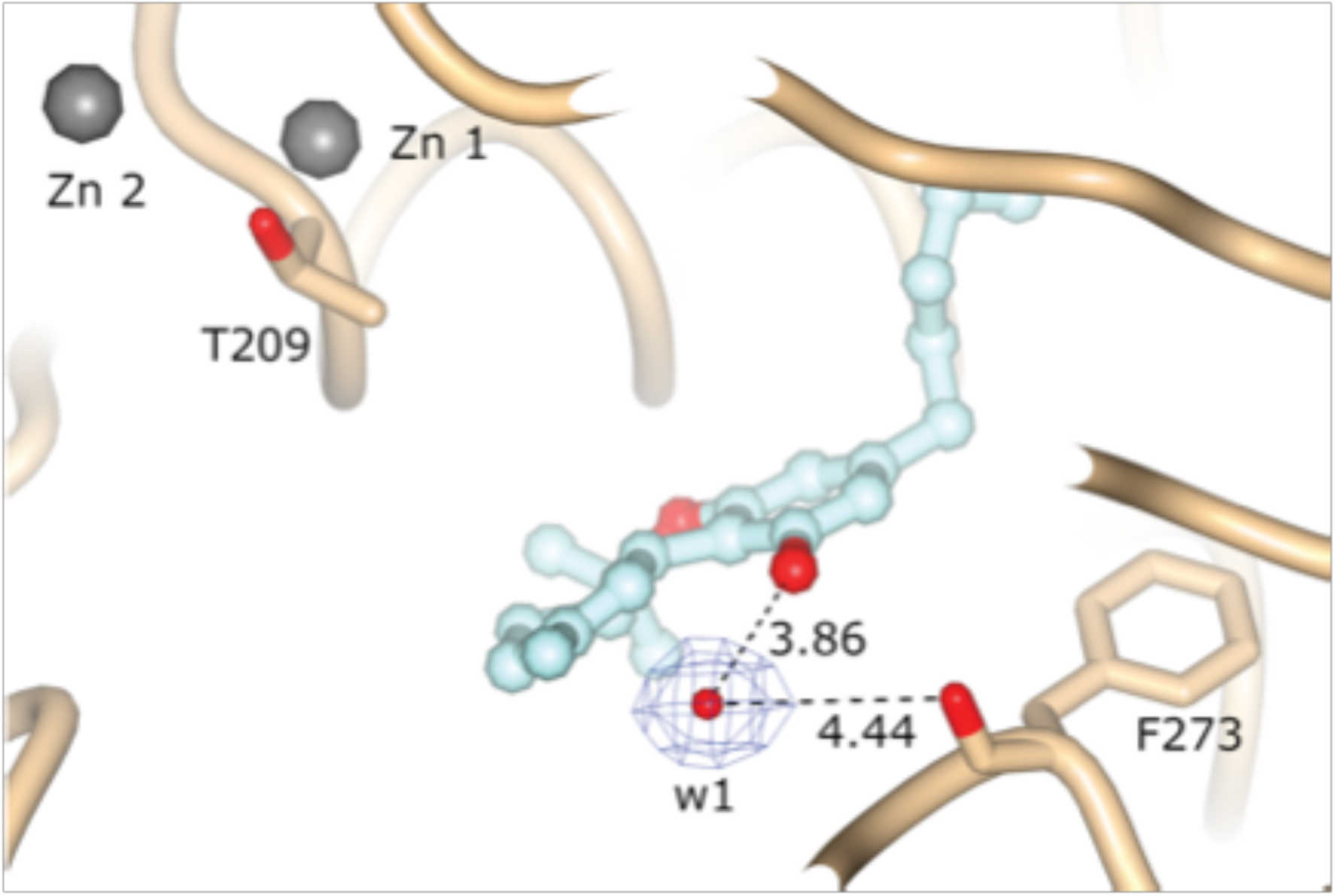
Water molecule present in the ATX-THC binding site. The distances from the water molecule to the oxygen of THC and carbonyl oxygen of F273 are indicated in Å.

**Fig. EV4.**
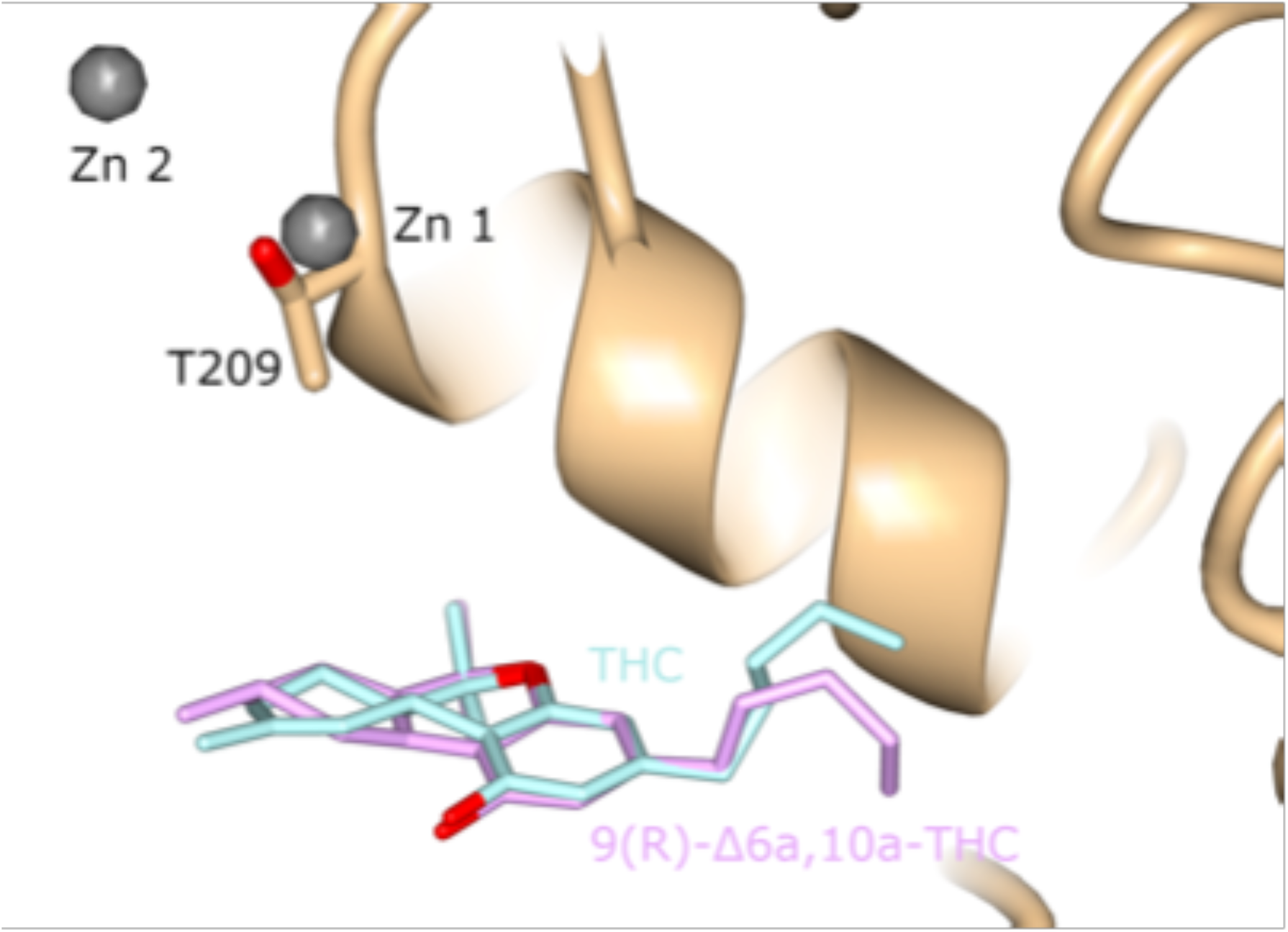
Superposition of the cannabinoids from ATX-THC and ATX-6a10aTHC. THC and 6a10aTHC are colored pink and turquoise, respectively.

**Fig. EV5.**
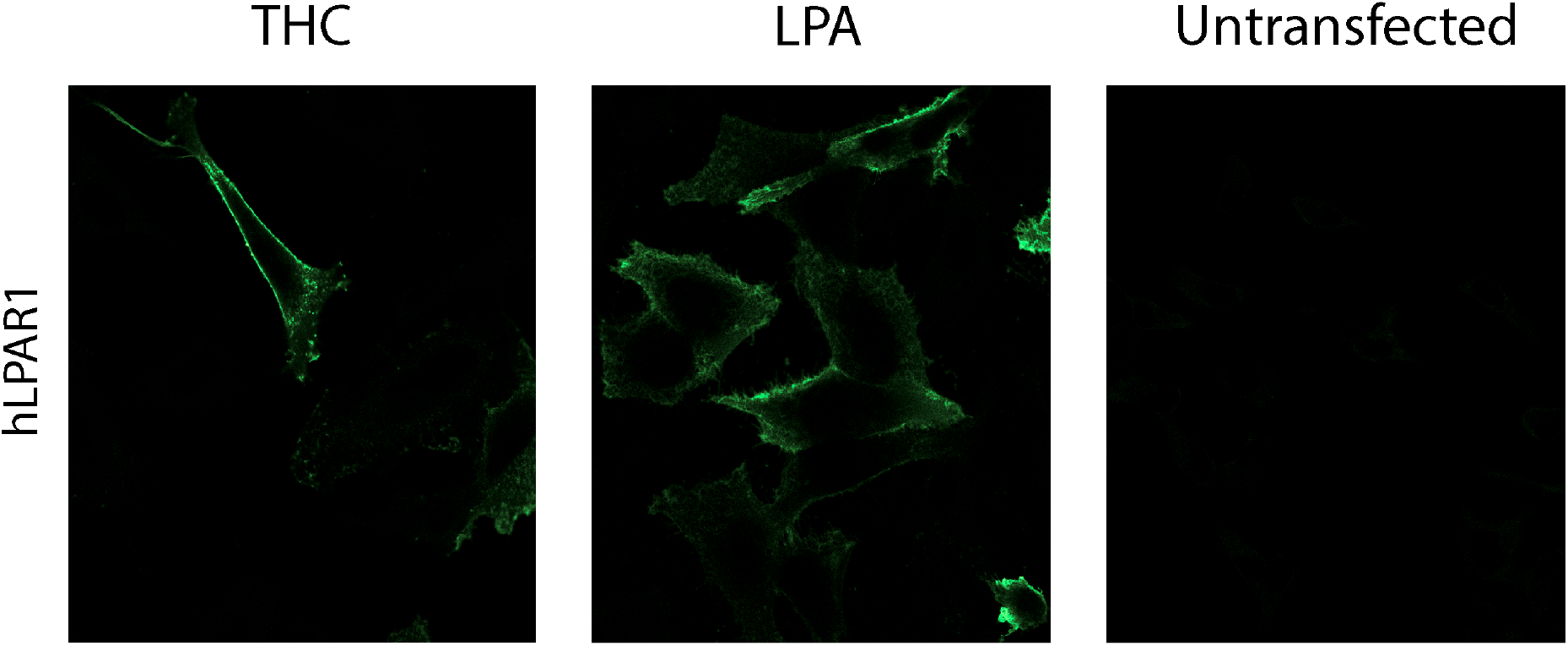
Control assays for LPA_1_ receptor internalization. No internalization was found using THC alone. LPA treatment of cells led to high LPA_1_ receptor internalization, in accordance with previously published results. Untransfected cells were not stained using primary and secondary antibody, confirming the specificity of the labelling reagents used.

**Fig. EV6.**
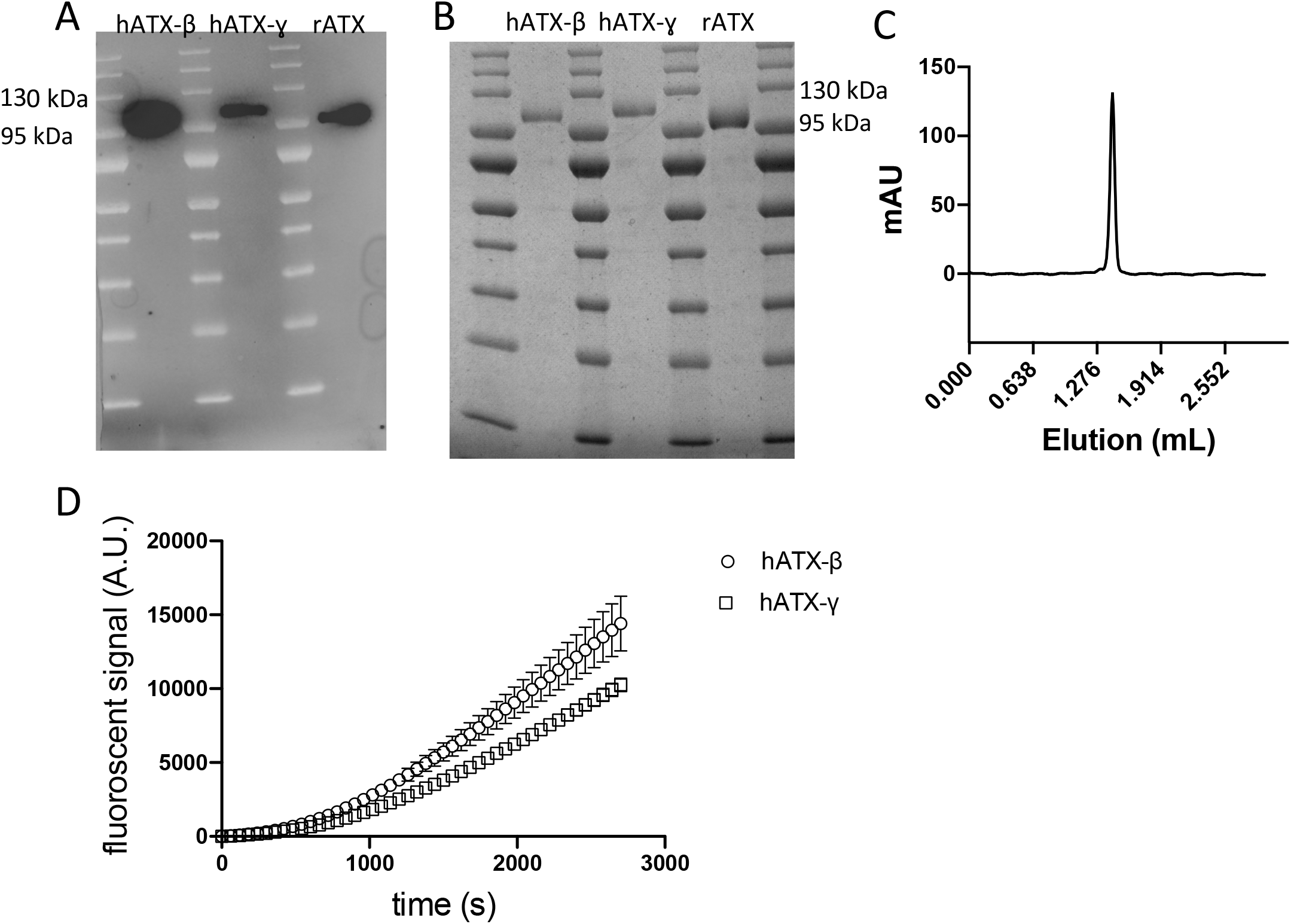
Recombinant ATX protein purity and enzymatic assays (A) Western blot analysis of recombinant proteins hATX-β, hATX-γ and rATX. (B) SDS-page analysis of recombinant proteins hATX-β, hATX-γ and rATX showing high purity. (C) Analytical gel filtration chromatography profile of rATX (D) Enzymatic curves superposition of hATX-β and hATX-γ.

